# Epigenetic signatures associated with imprinted paternally-expressed genes in the *Arabidopsis* endosperm

**DOI:** 10.1101/423137

**Authors:** Jordi Moreno-Romero, Gerardo Del Toro-De León, Vikash Kumar Yadav, Juan Santos-González, Claudia Köhler

**Affiliations:** Department of Plant Biology, Uppsala BioCenter, Swedish University of Agricultural Sciences and Linnean Center for Plant Biology, Uppsala, Sweden

**Keywords:** H3K9me2, H3K27me3, CHG methylation, endosperm, genomic imprinting, paternally expressed genes

## Abstract

**Background:** Imprinted genes are epigenetically modified during gametogenesis and maintain the established epigenetic signatures after fertilization, causing parental-specific gene expression.

**Results:** In this study, we show that imprinted paternally-expressed genes (PEGs) in the *Arabidopsis* endosperm are marked by an epigenetic signature of Polycomb Repressive Complex2 (PRC2)-mediated H3K27me3 together with heterochromatic H3K9me2 and CHG methylation, which specifically mark the silenced maternal alleles of PEGs. The co-occurrence of H3K27me3 and H3K9me2 on defined loci in the endosperm drastically differs from the strict separation of both pathways in vegetative tissues, revealing tissue-specific employment of repressive epigenetic pathways in plants. Based on the presence of this epigenetic signature on maternal alleles we were able to predict known PEGs at high accuracy and identified several new PEGs that we confirmed using INTACT-based transcriptomes generated in this study.

**Conclusions:** The presence of the three repressive epigenetic marks, H3K27me3, H3K9me2, and CHG methylation on the maternal alleles in the endosperm serves as a specific epigenetic signature that allows to predict genes with parental-specific gene expression. Our study reveals that there are substantially more PEGs than previously identified, indicating that paternal-specific gene expression is of higher functional relevance than currently estimated. The combined activity of PRC2-mediated H3K27me3 together with the heterochromatic H3K9me3 has also been reported to silence the maternal *Xist* locus in mammalian preimplantation embryos, suggesting convergent employment of both pathways during the evolution of genomic imprinting.

## BACKGROUND

Genomic imprinting is an epigenetic phenomenon causing maternal and paternal alleles to be differentially expressed after fertilization. In plants, genomic imprinting is mainly confined to the endosperm, an ephemeral nutritive tissue supporting embryo growth, similar to the placenta in mammals [1]. The endosperm is the product of a double fertilization event, where one of the haploid sperm cells fertilizes the haploid egg cell giving rise to the diploid embryo, while the second sperm cell fertilizes the diploid central cell to give rise to the triploid endosperm [2]. Imprinted genes are epigenetically modified during gamete formation and the established epigenetic asymmetry is maintained after fertilization. Differential DNA methylation is established by the DNA glycosylase DEMETER (DME) that removes methylated cytosine residues in the central cell of the female gametophyte [3]. DME is not active in sperm, leading to differential DNA methylation between male and female genomes in the endosperm. DME acts on small transposable elements (TEs) in the vicinity of genes [4] and its activity has been connected to the expression of maternally-expressed imprinted genes (MEGs). Hypomethylation can furthermore expose binding sites for the Fertilization Independent Seed (FIS)-Polycomb Repressive Complex 2 (PRC2) [5, 6], an evolutionary conserved chromatin modifying complex that applies a trimethylation mark on histone H3 at lysine 27 (H3K27me3) [7]. The Arabidopsis FIS-PRC2 consists of the subunits MEDEA (MEA), FIS2, FERTILIZATION INDEPENDENT ENDOSPERM (FIE), and MULTICOPY SUPPRESSOR OF IRA1 (MSI1) [8] and is specifically active in the central cell of the female gametophyte and in the endosperm [9]. Repression of the maternal alleles of PEGs is mediated by the activity of the FIS-PRC2 [10], consistent with maternal PEG alleles being marked by H3K27me3 [6]. In this manuscript, we addressed the mechanism of maternal allele repression in PEGs. We surprisingly found that PEGs are regulated by two otherwise largely exclusive epigenetic repressive pathways, the FIS2-PRC2 and the pathway establishing the heterochromatin-localized H3K9me2 modification [11]. We demonstrate that both modifications, H3K27me3 and H3K9me2, overlap on the maternal alleles of the majority of PEGs. Our data suggest that most likely FIS-PRC2 acts first and is required to establish H3K9me2. Furthermore, we find maternal alleles of PEGs to be marked by CHG methylation in the central cell, indicating that repressive pathways establishing H3K27me3, H3K9me2 and CHG methylation act in the central cell of the female gametophyte. Finally, we use the presence of the three modifications to predict novel PEGs and propose that the number of PEGs predicted based on expression data strongly underestimates the real number of PEGs.

## RESULTS

### The maternal alleles of PEGs are marked by H3K27me3, H3K9me2, and CHG methylation

In a previous study, we revealed that H3K27me3 and H3K9me2 overlapped at paternal heterochromatic regions in the endosperm, suggesting a partial functional redundancy of both modifications [6]. We now addressed the question whether this redundancy extends to other regions of the genome. We found that about one-third of genes marked by H3K27me3 on the maternal alleles were also marked by H3K9me2 (Fig. 1a, hypergeometric test, *P*=0; Additional file 1: Figure S1A, *P*=0). Genes containing both modifications on their maternal alleles had significantly higher levels of H3K27me3 compared to those only marked by H3K27me3 (Fig. 1b, Additional file 1: Figure S1B). The majority of double-marked genes contained both modifications specifically on the maternal but not the paternal alleles (Fig. 1a, Additional file 1: Figure S1A) and increasing levels of H3K27me3 on maternal alleles correlated with increasing levels of H3K9me2 on the maternal but not the paternal alleles (Additional file 1: Figure S2 and S3). We thus proposed that the presence of both modifications on the maternal alleles correlates with paternally-biased expression. Consistent with this notion we found that genes previously identified as PEGs [12] were substantially enriched for both modifications on their maternal alleles, while MEGs did not show enrichment of both marks on either maternal or paternal alleles (Fig. 1c and 1d, Additional file 1: Figure S1C-D). This trend was independent of the direction of the cross and similarly observed in Col x L*er* and L*er* x Col crosses (Fig. 1 and Additional file 1: Figure S1). The presence of H3K27me3 is generally confined to gene bodies [13], consistent with the observed enrichment of H3K27me3 in gene bodies of PEG maternal alleles (Fig. 1e and Additional file 1: Figure S1E). Interestingly, H3K9me2 was similarly restricted to gene bodies of PEGs (Fig. 1e and Additional file 1: Figure S1E), contrasting the exclusion of H3K9me2 from genic regions in sporophytic tissues [14]. The maternal alleles of PEGs were also strongly enriched for CHG methylation (Fig. 1f, Additional file 1: Figure S1 and S2), consistent with the chromomethylase 3 (CMT3) acting in a positive feedback loop with the histone methyltransferase proteins KYP/SUVH4, SUVH5, and SUVH6 [11]. Levels of CHH methylation were generally higher on maternal than on paternal alleles (Fig. 1f, Additional file 1: Figure S1F), consistent with depletion of CHH methylation in sperm [4, 15].

**Fig. 1.**
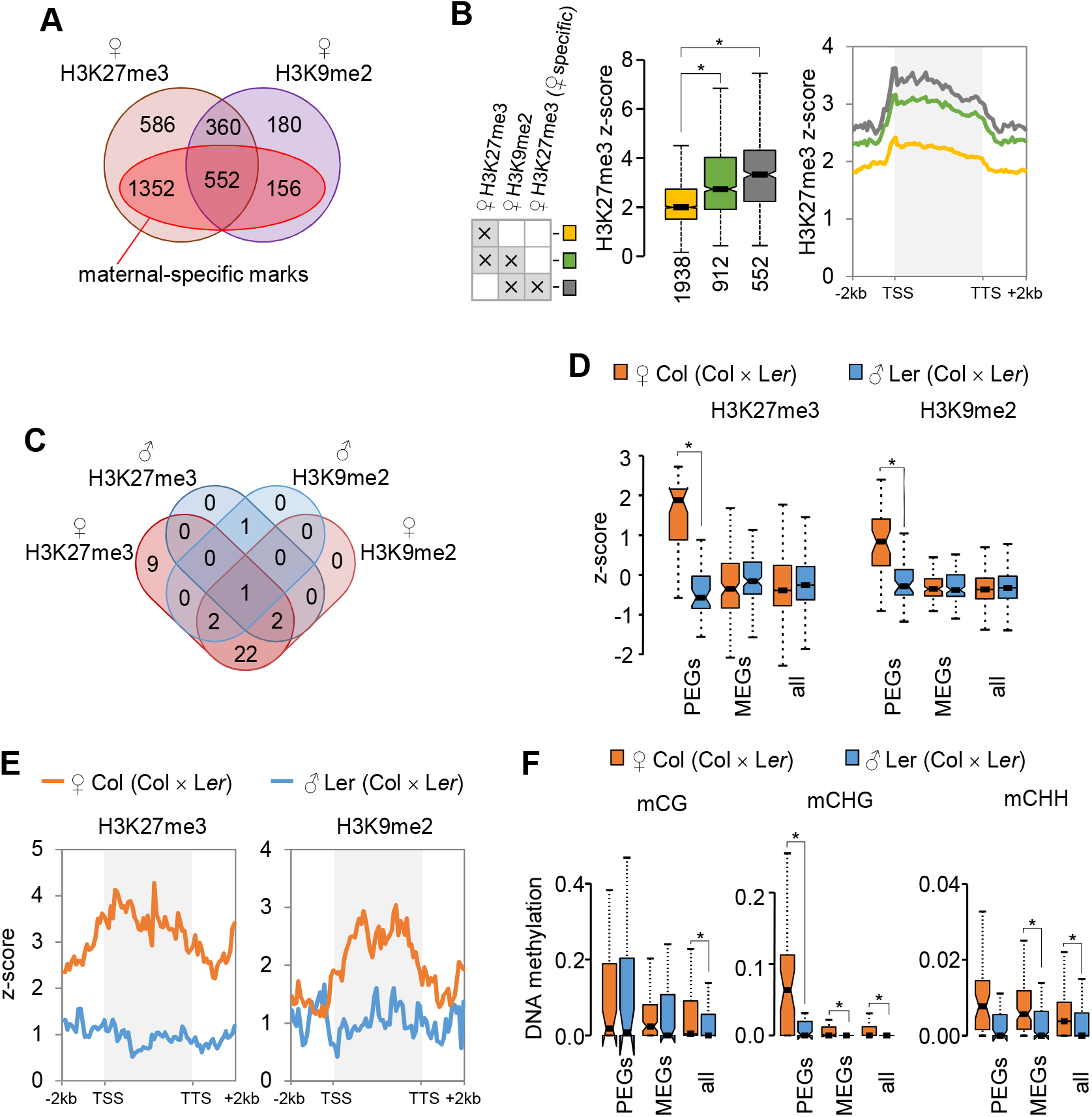
H3K27me3, H3K9me2, and CHG methylation overlap on maternal alleles of PEGs. **a** Overlap of genes marked by H3K27me3 and H3K9me2 on the maternal alleles. Dark red inset marks genes that have H3K27me3 and H3K9me2 exclusively on their maternal alleles (excluding genes also marked on the paternal alleles). **b** Genes marked by H3K27me3 and H3K9me2 have higher levels of H3K27me3. Box plots show mean values of z-scored H3K27me3 and H3K9me2 of genes where the maternal alleles are either marked by H3K27me3 alone or by a combination of H3K27me3 and H3K9me2. Genes containing maternal-specific H3K27me3 and H3K9me2 are particularly enriched for H3K27me3. Boxes show medians and the interquartile range, error bars show the full range excluding outliers. Notches show the 95% confidence interval for the median. Asterisks mark significance (Wilcoxon rank sum test, *P* <1.0E-3). Presence of the mark corresponds to the mark highest score defined in Additional File 2: Table S1. **c** The majority of PEGs are marked by H3K27me3 and H3K9me2 on their maternal alleles. Overlap of PEGs [12] with maternal and paternal H3K27me3 and H3K9me2. **d** Levels of H3K27me3 and H3K9me2 on PEGs, MEGs, and all genes. Box plots (as specified in panel B) show mean values of z-scored H3K27me3 and H3K9me2. Asterisks mark significance (Wilcoxon rank sum test, *P* <1.0E-3). **e** Metagene plots showing average distribution of z-score normalized H3K27me3 and H3K9me2 levels on maternal and paternal alleles of PEGs [12]. **f** Levels of DNA methylation in each sequence context (mCG, mCHG and mCHH) on PEGs, MEGs, and all genes. Box plots (as specified in panel B) show mean values of relative DNA methylation. Asterisks mark significance (Wilcoxon rank sum test, *P*<5.0E-3). Data shown correspond to cross Col × L*er* [6], for reciprocal cross direction see Additional file 1: Figure S1.

### Maternal-specific CHG methylation is established in the central cell and depends on FIS-PRC2

PEGs had increased levels of CHG methylation in the central cell, but low CHG methylation in sperm and vegetative cells of pollen (Fig. 2a), revealing that differences in CHG methylation are established before fertilization. The FIS-PRC2 had been proposed to promote non-CG methylation [4], raising the hypothesis that increased CHG methylation on the maternal alleles of PEGs depends on the activity of FIS-PRC2 in the central cell. Consistently, we observed strongly decreased CHG methylation on the maternal alleles of PEGs in seeds lacking maternal FIE activity (Fig. 2b), suggesting that FIS-PRC2 activity is required to recruit CHG methylation. The DNA glycosylase DME is required for the activation of *MEA* and *FIS2*, both encoding subunits of the FIS-PRC2 [16, 17]. Maternal PEG alleles had reduced levels of CHG methylation in seeds lacking maternal DME activity (Fig. 2b), supporting the notion that FIS-PRC2 activity is required for CHG methylation establishment on maternal PEG alleles.

**Fig. 2.**
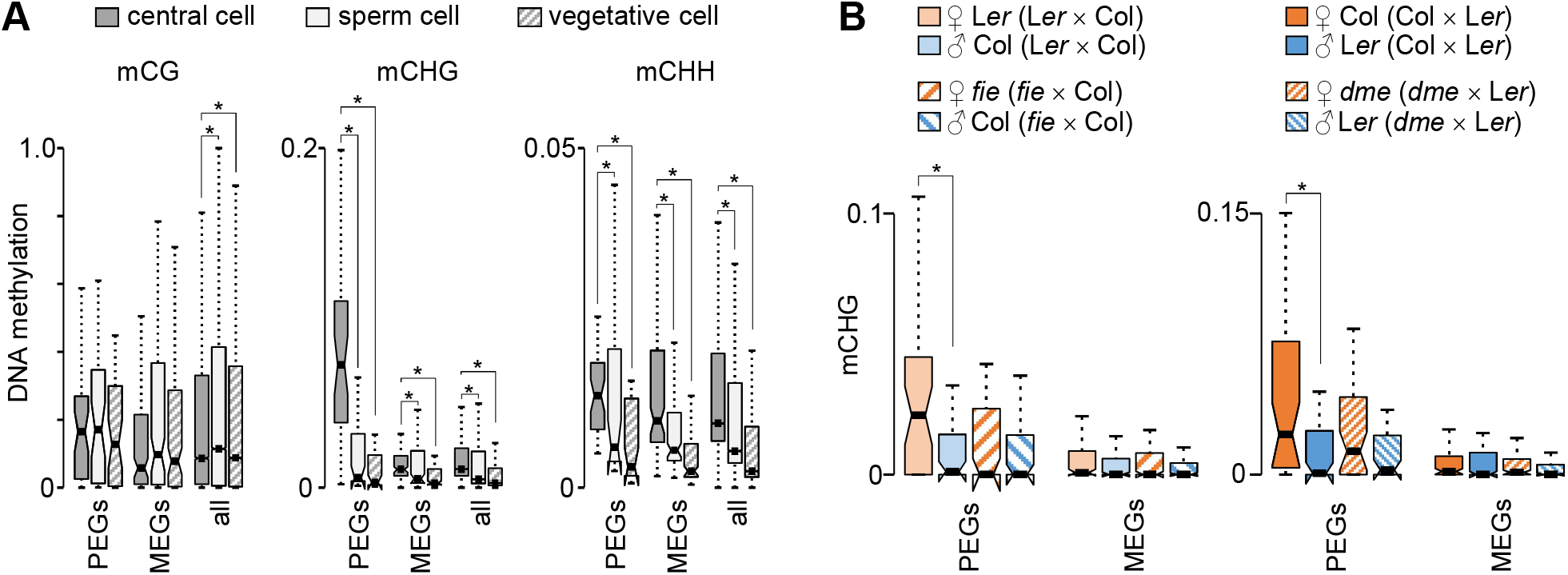
Maternal CHG methylation is established in the central cell and depends on FIS-PRC2. **a** Box plots showing mean values of relative DNA methylation in the central cell, sperm cell, and vegetative cell for PEGs, MEGs, and all genes. Boxes show medians and the interquartile range, error bars show the full range excluding outliers. Notches show the 95% confidence interval for the median. Asterisk marks significance (Wilcoxon rank sum test, *P* <5.0E-3). **b** Box plots showing mean values of CHG methylation for PEGs and MEGs on maternal and paternal alleles in wild-type, *fie* (left panels) and *dme* (right panels) endosperm. Asterisk marks significance (Wilcoxon rank sum test, *P* <5.0E-2).

Previous work revealed that loss of FIS-PRC2 function caused activation of maternal PEG alleles [10]. We addressed the question whether loss of CMT3 and the redundantly acting SUVH4,5,6 [18] similarly affect silencing of maternal PEG alleles. Based on published expression studies, *CMT3*, *SUVH4*, and *SUVH6* are expressed in the central cell of the female gametophyte [19]. The maternal alleles of seven tested PEGs remained silenced in reciprocal crosses of wild type with *cmt3* and *suvh4,5,6* triple mutants (Additional file 1: Figure S4A-B), indicating that stable silencing of the maternal alleles of PEGs does not depend on CMT3 and SUVH4,5,6 activity before fertilization. We tested the requirement of CMT3 and SUVH4,5,6 for PEG regulation after fertilization by monitoring PEG expression in homozygous mutant *cmt3* and *suvh4,5,6* seeds. None of the seven tested genes was significantly upregulated in seeds of *cmt3* or *suvh4,5,6* mutants (Additional file 1: Figure S4C), indicating that CMT3 and SUVH4,5,6 are not required for repression of maternal PEG alleles after fertilization. In the *suvh4,5,6* triple mutant, H3K9me2 is genome-wide eliminated in vegetative tissues [20]; however, whether H3K9me2 is similarly eliminated in the central cell and endosperm remains to be shown. Therefore, deciphering the role of H3K9me2 in repressing the maternal alleles of PEGs remains to be subject of future investigations.

### Paternally-biased expression coincides with the combination of H3K27me3, H3K9me2, and CHG methylation

We addressed the question whether the presence of H3K9me2 and CHG methylation is functionally relevant for maternal allele repression by testing whether the presence of both modifications either alone or in combination with H3K27me3 correlates with maternal allele repression. Strikingly, while the presence of either CHG methylation, H3K9me2, or H3K27me3 on maternal alleles did not shift the allelic balance, the combination of more than one modification shifted the balance towards preferential paternal allele expression (Fig. 3a, Additional file 1: Figure S5A), suggesting that the combined presence of H3K27me3, H3K9me2, and CHG methylation is a hallmark for maternal allele repression. Consistently, increased bias towards paternal allele expression correlated with increased levels of H3K27me3, H3K9me2, and CHG methylation on maternal alleles (Fig. 3b, Additional file 1: Figure S5B).

**Fig. 3.**
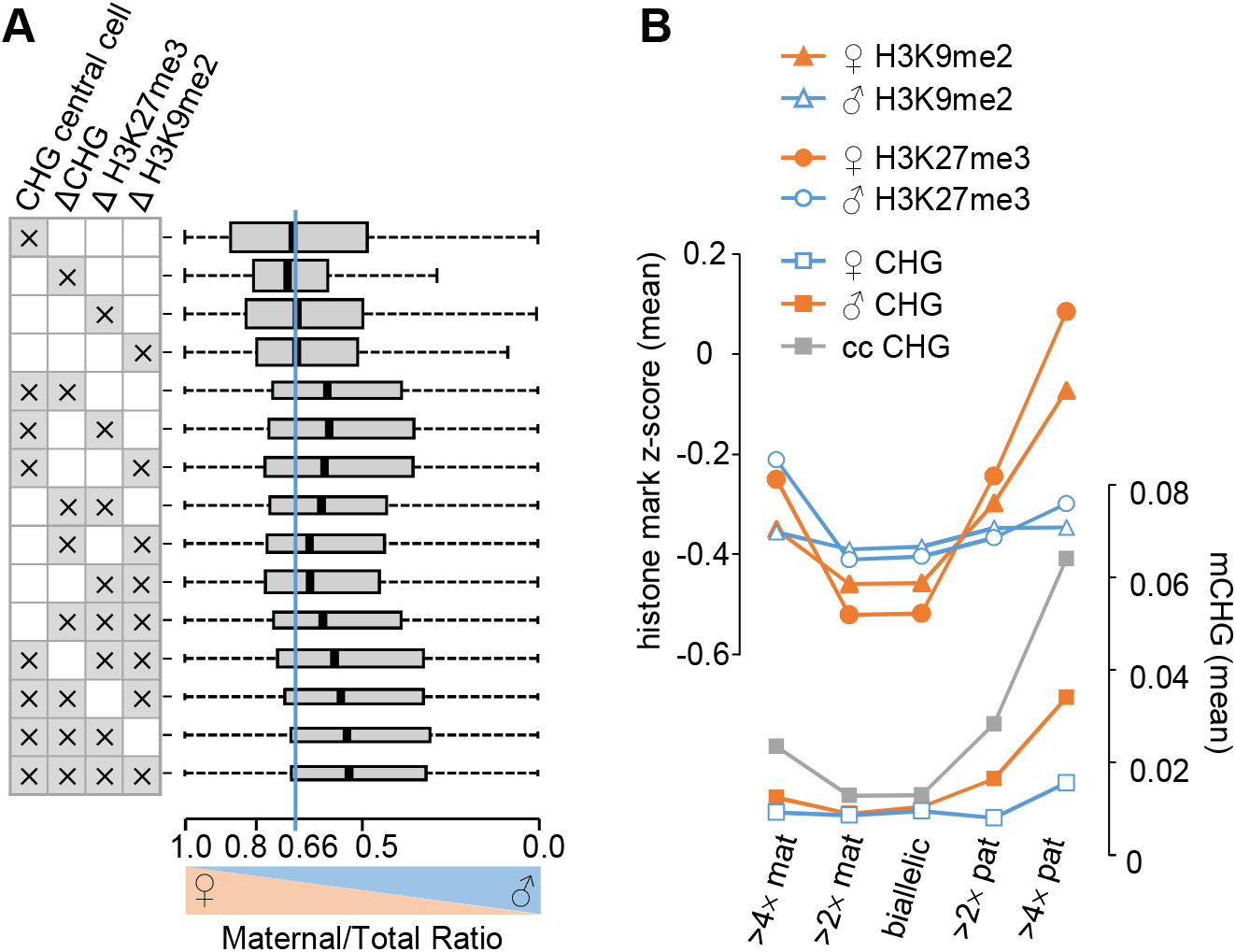
Paternally-biased expression coincides with the combination of H3K27me3, H3K9me2, and CHG methylation. **a** Box plots show mean values of maternal to total reads for genes marked by one or several of the following modifications: CHG methylation in the central cell, difference of CHG methylation, H3K27me3, or H3K9me2 levels on maternal and paternal alleles in the endosperm. For each modification only genes present in the highest scoring category were included in the analysis (see Additional File 2: Table S1). Boxes show medians and the interquartile range, error bars show the full range excluding outliers. Widths of boxes are proportional to square-root of the number of genes in each category. Blue line indicates the expected ratio for biallelically expressed genes. **b** Paternal expression bias correlates with high levels of CHG methylation in the central cell and H3K27me3 and H3K9me2 on maternal alleles in the endosperm. Plotted are mean values of CHG methylation, H3K27me3, and H3K9me2 on maternal and paternal alleles for genes with deviating parental expression levels from the expected two maternal to one paternal ratio. Data shown correspond to cross Col × L*er*, for reciprocal cross direction see Additional file 1: Figure S5.

We tested which combination of marks would most reliably allow predicting paternally-biased genes. We assigned scores for the allele-specific presence of the three modifications (see Additional file 2: Table S1) and tested whether highly scoring genes were enriched for previously predicted PEGs [12]. The combination of CHG methylation in the central cell together with maternal-specific H3K27me3 and H3K9me2 allowed to predict the highest number of previously described PEGs in Col and L*er* accessions (24 out of 42 (57%), [12]) in relation to the number of genes in the category with the highest score (Fig. 4a, Additional file 2: Table S1) and was chosen for further analysis. The category with the highest score was significantly enriched for PEGs (hypergeometric test, *P*=1.0e-32); while categories with lower scores contained only few PEGs (Fig. 4a). Similarly, out of 64 PEGs that had been predicted by a recent study re-evaluating previously published imprintome datasets [21], 40 (62%) PEGs were present in the highest score category (Fig. 4a). Nearly half (96 genes, 46.4%) of those genes in the highest score category were significantly paternally biased (Chi-square <0.05, Bonferroni corrected, Additional file 3: Table S2), which was significantly more than the 8% paternally-biased genes identified among all genes tested (Hypergeometric test, *P*= 8.9e-52). We thus conclude that the presence of the three modifications, CHG in the central cell, H3K27me3 and H3K9me2 on maternal alleles in the endosperm allows to predict genes with paternally-biased expression. Paternally biased genes were particularly involved in transcriptional regulation (*P*=7.54e-5) and chromatin organization (*P*=1.57e-3), consistent with previous reports on the functional role of PEGs [12, 22].

**Fig. 4.**
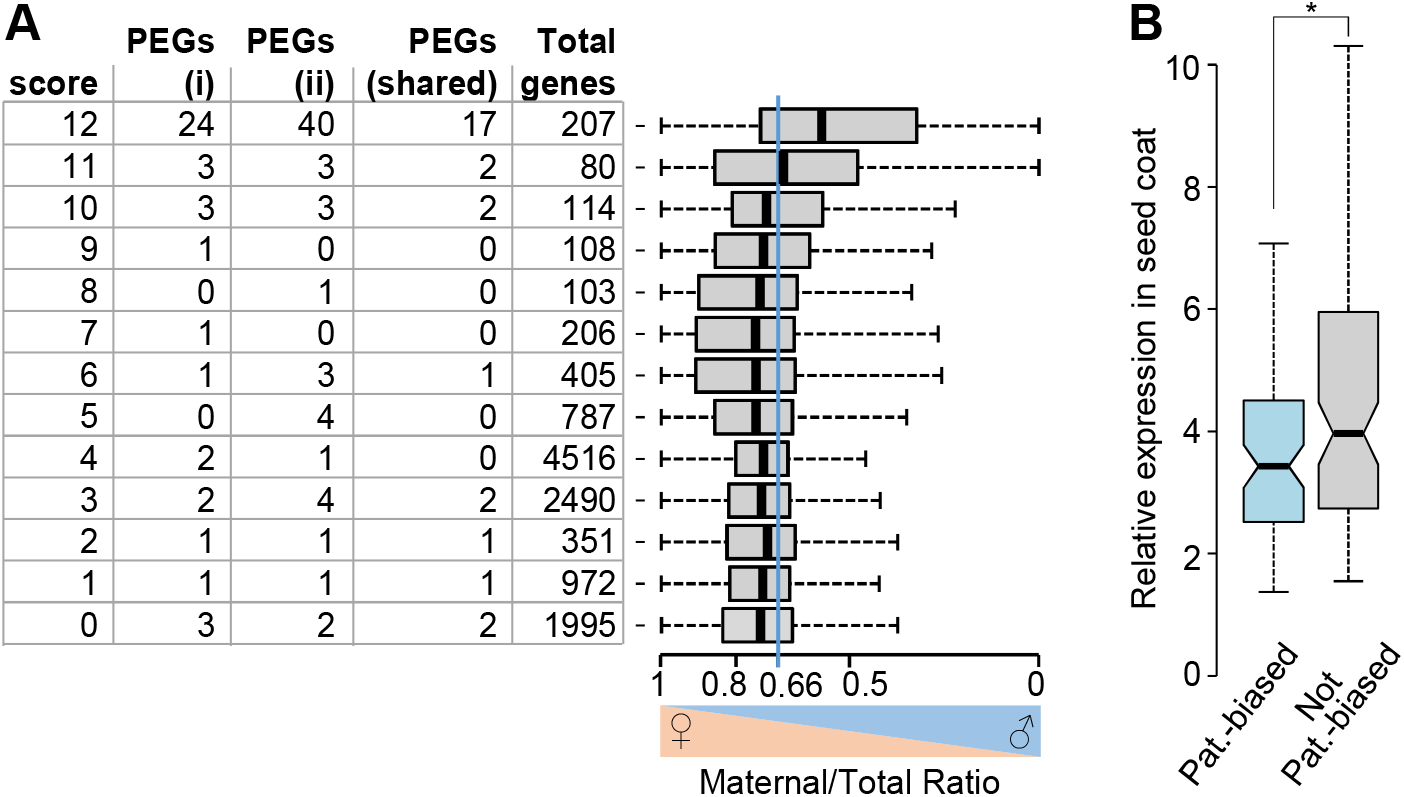
Epigenetic signatures allow predicting paternally-expressed imprinted genes (PEGs). **a** The combination of CHG methylation in the central cell together with maternal-specific H3K27me3 and H3K9me2 allows predicting previously described PEGs [12, 21]. Scores are calculated as described in the Additional File 2: Table S1. High and low scores correspond to high or low levels of CHG methylation, H3K27me3, and H3K9me2, respectively. Number of PEGs corresponds to PEGs previously predicted [12] (i) and [21] (ii) in Col and L*er* accessions. Total genes correspond to all genes present in the corresponding categories. Box plots show mean values of maternal to total reads in the indicated categories. Boxes show medians and the interquartile range, error bars show the full range excluding outliers. Blue line indicates the expected ratio for biallelically expressed genes.**b** Box plots show relative expression level of genes in the seed coat of torpedo stage embryos [23]. Genes present in the highest score category (Additional File 3: Table S2) were grouped in paternally biased genes (Chi-square test, *P*<0.05 Bonferroni corrected) and not-paternally biased genes. Asterisks mark significance (Wilcoxon rank sum test, *P* <5.0E-3).

### Maternal seed coat contamination restricts the identification of PEGs

Published endosperm transcriptome data contain a substantial fraction of transcripts from the maternal seed coat, which may limit the correct prediction of paternally-biased genes [21]. We hypothesized that there are several genes that based on their epigenetic modifications (score 12, Fig. 4a and Additional file 3: Table S2) are likely to be PEGs but based on available expression data are not correctly classified. A prediction of this hypothesis is that genes in the highest score category that have been classified as biallelically expressed or maternally-biased are more highly expressed in the seed coat than those with paternally-biased expression. We tested this hypothesis using available expression data of seed coat tissue [23] at a similar developmental stage as published endosperm expression data [12]. Indeed, genes with paternally-biased expression were significantly less expressed in the seed coat compared to genes that based on available expression data are either predicted to be biallelically expressed or maternally-biased (Wilcoxon rank sum test, *P*<0.005; Fig. 4b), strongly suggesting that maternal tissue contamination in available expression data limits the prediction of PEGs. To further test this hypothesis, we generated transgenic reporter lines containing the promoter (≃ 2kb) and downstream genic regions fused to the green fluorescent protein (GFP) of seven genes belonging to the highest score category but predicted to be maternally (*AT2G33620*, *AT1G43580*, *AT1G47530*, *AT1G64660*, *AT2G30590*, *AT4G15390*) or biallelically expressed (*AT5G53160*). We detected a GFP signal in the endosperm only for construct *AT1G64660* (Fig. 5, Additional file 1: Table S3). For construct *AT1G47530* a signal was detected in the seed coat, while for the other constructs no GFP signal was detected in seeds, indicating that the regulatory elements required for the expression of those genes are located outside the promoter and genic regions used to generate the reporter lines. Reciprocal crosses using the *AT1G64660* reporter lines revealed that this gene is indeed a PEG and strongly expressed in the endosperm when paternally, but not when maternally inherited (Fig. 5). This data support the hypothesis that seed coat contamination limits the transcriptome-based identification of PEGs.

**Fig. 5.**
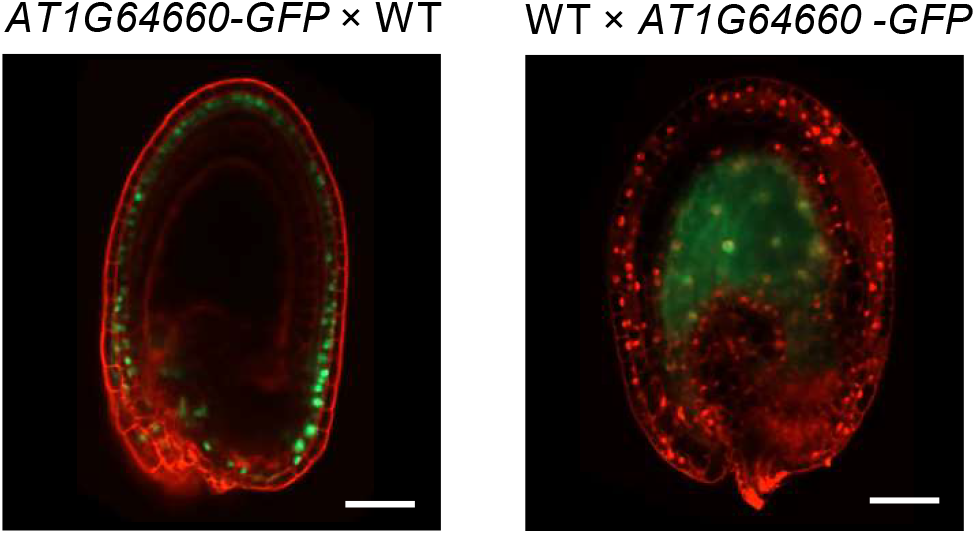
Reporter lines for *AT1G64660* confirm imprinted expression. Representative images of seeds derived after reciprocal crosses of the *AT1G64660* reporter line (fusion with green fluorescent protein (GFP)) with wild-type (WT) plants. GFP fluorescence was detected in the seed coat when the *AT1G64660* reporter was maternally inherited, but endosperm-specific expression was only detected when the *AT1G64660* reporter was paternally inherited. Seeds at 2 DAP were used for imaging. Red stain is propidium iodide. Scale bars correspond to 50 μm.

To test whether we could confirm additional PEGs that have been predicted based on their epigenetic signatures, we applied INTACT (isolation of nuclei tagged in specific cell types) to purify endosperm nuclei at 4 days after pollination (DAP) from Col × L*er* and L*er* × Col reciprocal crosses. Isolated RNA was sequenced and profiled for allele-specific gene expression. By analyzing the maternal to total reads ratio in each epigenetic category, we confirmed that the genes in the highest score category (group with score 12) indeed showed a clear trend towards paternally-biased expression (Fig. 6a, Additional file 4: Table S4). Following previously established criteria [12], we predicted 148 PEGs that were reciprocally imprinted in both directions of the crosses. There was a significantly higher number of PEGs present in the highest score category compared to a representative random sample of genes with informative reads (Fig. 6b). Furthermore, the highest score category had the highest frequency of PEGs, with other categories having significantly fewer PEGs (Fig. 6c). Of the 148 genes that we predicted as PEGs based on our RNA sequencing data, 45 were present in the highest score category, which is significantly more than expected by chance (p=2.008 e-55, Fig. 6d). Out of those, 24 were previously predicted based on published data [10, 12, 24, 25], while 21 genes are likely new high-confidence PEGs, revealing that PEGs are more common than previously estimated.

**Fig. 6.**
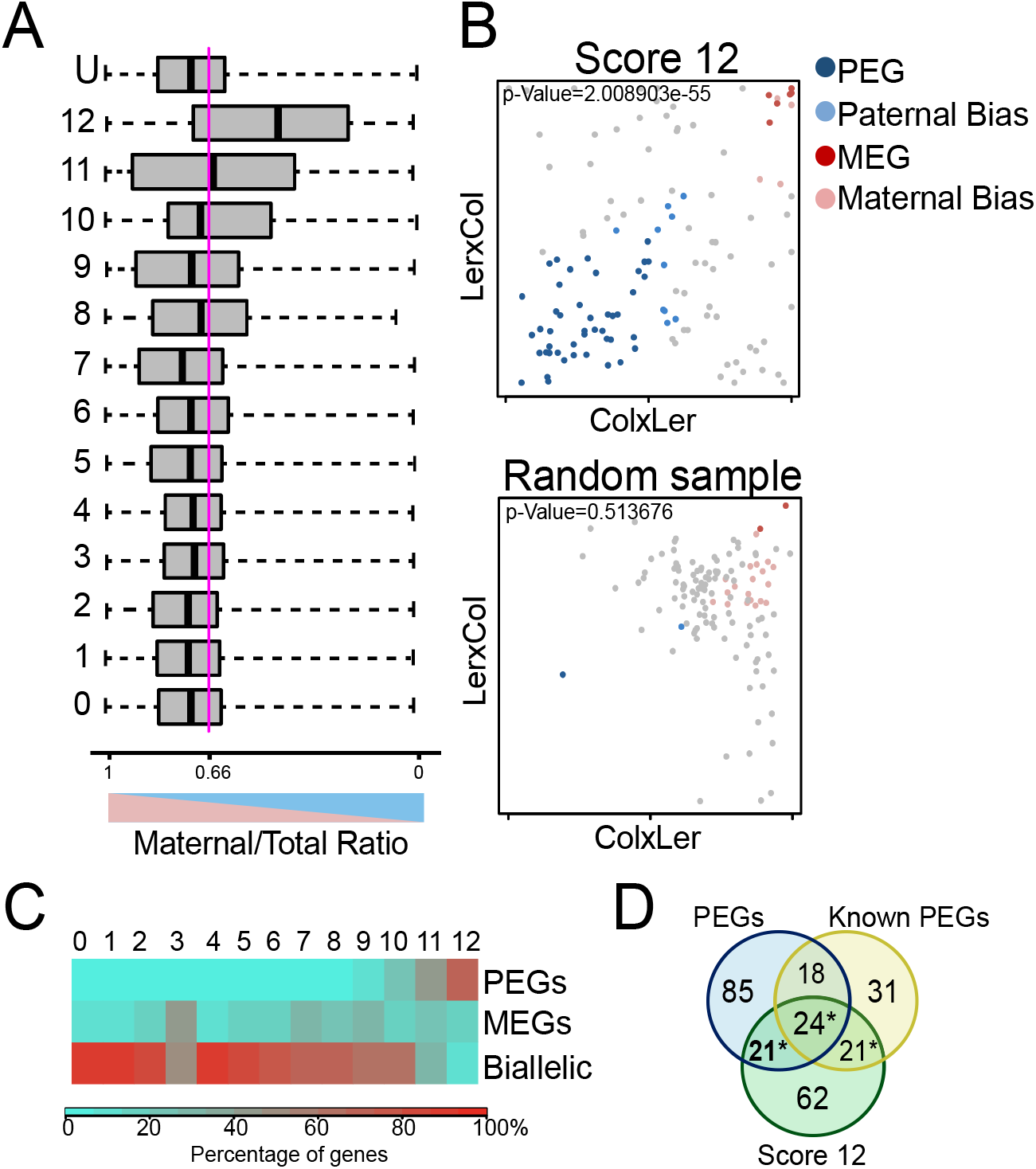
Confirmation of predicted PEGs in RNA samples of INTACT-purified endosperm nuclei. **a** Distribution of maternal to total read values for genes in each epigenetic category. The distribution for the total population of genes with informative reads (U) is included. Genes with the highest levels of CHG methylation, H3K27me3, and H3K9me2 show a clear trend towards paternally-biased expression (around 0.5). Values correspond to combined values of Col × L*er* and L*er* × Col crosses. Boxes show medians and the interquartile range, lines show the full range of outliers. Blue line indicates the expected ratio for biallelically expressed genes (0.66). **b** Parent-of-origin distribution of the group of genes with score 12, compared with a random group of the same size. Significantly more PEGs are detected in the group of genes with score 12, compared with the random sample (Hypergeometric test, *P*-values correspond to the comparison with the total population of PEGs in the whole RNAseq data). The scatter plots show maternal to total read values for Col × L*er* and L*er* × Col reciprocal crosses. Colors indicate genes with significant parent-of-origin biased expressed: MEGs, 85% or more of maternal reads in each reciprocal cross; PEGs, 50% or more of paternal reads in each reciprocal cross. Maternally and paternally biased genes show significantly biased expression (*P* <= 0.01) in both reciprocal crosses. **c** Percentage of PEGs, MEGs and biallelically expressed genes in each epigenetic category. Color code indicates the percentage of genes within each category. Categories with genes having lower levels of CHG methylation, H3K27me3, and H3K9me2 marks have significantly fewer PEGs than the highest scoring category (Chi-square test, *P*<1e-8 in categories 1-10; *P*=0.001 in category 11). Only PEGs, MEGs and Biallelic categories were considered. **d** Overlap of PEGs that were detected in our INTACT-based RNAseq samples (blue) with genes predicted as PEGs based on their epigenetic score 12 (green) and previously reported PEGs (yellow) by [10, 12, 24, 25]. New predicted and confirmed PEGs in bold. Statistical significance was calculated using a hypergeometric test (*P*-values<4e-20) and significant overlaps are marked by asterisks.

## DISCUSSION

In this study, we identified the concomitant presence of maternal-specific CHG methylation, H3K27me3, and H3K9me2 as an epigenetic signature for paternally-biased expression in the endosperm. We furthermore predict that there are substantially more PEGs than previously reported in Arabidopsis [10, 12, 24], suggesting that in Arabidopsis the number of PEGs exceeds the number of MEGs. Recent re-evaluation of published imprintome data of Arabidopsis revealed that a large number of previously predicted MEGs were seed coat expressed genes, while the number of PEGs was underestimated [21]. Our study supports and extends this notion by showing that there is a large number of paternally-biased genes that likely failed to be identified in previous studies because of maternal seed coat contamination or early stage-specific expression.

A previous study reported that the maternal alleles of PEGs in *A. lyrata* are marked by CHG methylation and implicated that closely related species use different mechanisms to regulate imprinted gene expression [26]. Our study reveals that similar to *A. lyrata* the maternal alleles of many PEGs are also marked by CHG methylation in *A. thaliana*, highlighting that epigenetic mechanisms employed to maintain monoallelic expression are rather conserved between related species. Interestingly, while the maternal alleles of PEGs in maize are marked by H3K27me3, they are not marked by CHG methylation [27], indicating that diverged species may use different mechanism in maintaining maternal allele repression.

How H3K9me2 and CHG methylation are established at PRC2 target genes remains to be studied; however, the strong activation of maternal PEG alleles upon loss of FIS-PRC2 function [10] suggests that H3K9me2 and CHG methylation require FIS-PRC2 function. The PRC2 is generally targeting genes with specific roles during development [28], while complexes establishing H3K9me2 are mainly targeting TEs localized in heterochromatic regions of the genome [29]. This functional division of PRC2 and machineries establishing heterochromatic marks is conserved in plants as well as in mammals [30]; however, there are notable exceptions to this rule in both groups of organisms. In rice seedlings, about one-third of H3K27me3 marked genes are also marked by CHG and CHH methylation [31]. Similar to the findings reported in our study, higher levels of H3K27me3 correlate with higher CHG and CHH methylation in gene bodies of rice [31]. The rice H3K27me3 methyltransferase SDG711 physically interacts with the CHH methyltransferase OsDMR2 and the SRA-domain containing SUVH protein SDG703, uncovering a mechanistic connection between PRC2 and non-CG methylation. In mammalian cells, H3K9 methyltransferases colocalize with PRC2 [32- 34], revealing crosstalk between these two major epigenetic silencing pathways that likely is required for stable gene silencing. The imprinted maternal *Xist* locus encoding an X-linked long-noncoding RNA is covered by H3K27me3 and H3K9me3 in preimplantation embryos [35, 36]. Importantly, loss of maternal H3K27me3 induces *Xist* activation, indicating that maternal H3K27me3 is the major imprinting mark of *Xist* [36]. This is strikingly similar to findings made in this study revealing H3K27me3 as the major repressive mark for PEGs. Recent work revealed that maternal H3K27me3 controls DNA methylation-independent imprinting in mammalian preimplantation embryos [37]. While imprinted expression of most genes is lost in the embryonic cell lineage, few genes maintain their imprinted expression in the extra-embryonic cell lineage [37]. The *Xist* locus that is marked by H3K27me3 and H3K9me3 is among those loci that remain imprinted in extra-embryonic tissues [36]. Whether the presence of both marks distinguishes those genes that maintain their imprinted expression from those that become biallelically expressed remains to be tested, but we consider this a very attractive hypothesis. We speculate that the presence of both marks in certain tissue types of mammals and flowering plants is a conserved epigenetic signature marking stably repressed genes.

## CONCLUSIONS

We discovered the co-occurrence of the PRC2-mediated H3K27me3 and heterochromatic H3K9me2 and CHG methylation as an epigenetic signature marking the silenced maternal alleles of PEGs. This signature can be used to predict PEGs at high accuracy and based on this prediction we estimate that the number of PEGs is substantially larger than previously estimated. We hypothesize that the common use of PRC2 and H3K9 methylation to silence target loci during reproduction has convergently evolved in flowering plants and mammals to ensure stable silencing during this sensitive life stage.

## METHODS

### Plant Material and Growth Conditions

All seeds were surface sterilized (5% sodium hypochlorite and 0.01% Triton X-100), stratified for 2 days at 4°C and germinated on half-strength Murashige and Skoog medium containing 1% sucrose under long-day conditions (16 h light/8 h darkness; 21°C). Plants were transferred to soil after 10 to 12 days and grown under long day conditions. The *cmt3-11* (SALK_148381; [38]) and *suvh456* mutants (kindly provided by Judith Bender) used in this study are in the Col-0 background.

### Imprinting Assays and Expression Analysis

To generate siliques of indicated crosses, three to five flowers were emasculated, hand-pollinated, and harvested at 4 DAP. RNA extraction was performed using the MagJET Plant RNA Purification Kit (Thermo Scientific) following the manufacturer’s instructions. Residual DNA was removed using Invitrogen DNase I (Amplification Grade), and cDNA was synthesized using the Fermentas first strand cDNA synthesis kit according to the manufacturer’s instructions. Quantitative PCR was performed using a MyiQ5 real-time PCR detection system (Bio-Rad) and Solis BioDyne-5x Hot FIREPol EvaGreen qPCR Mix Plus (ROX, Solis BioDyne). For the imprinting-by-sequencing assay, the PCR products were purified and analyzed by Sanger sequencing. For the imprinting-by-restriction enzyme digestion assay, the PCR products were purified and digested. Restriction enzymes and primers used are listed in the Additional file 1: Table S5.

### Data analysis

We made use of endosperm-specific ChIP-seq data that have been previously generated in our group [6]. Data correspond to pooled biological triplicates with ChIP signals being normalized with H3 ChIP data by calculating the log2 ratio in 150-bp bins across the genome. Data were standardized and normalized with a z-score transformation [39]. Metagene plots over genes were constructed between −2 kb and +2 kb by calculating mean levels of methylation signals in 100-bp bins in the flanks of the genes and in 40 equally long bins between the transcriptional start and stop. Gene z-scores were calculated as an average of z-scores over the gene body. DNA methylation data of *fie* and *dme* and their corresponding wild types are from [4]. DNA methylation data of the central cell, sperm, and vegetative cells are from [3] and [15]. Endosperm expression data are from [12] and seed coat expression data from [23].

### Endosperm nuclei isolation and RNA sequencing

We performed Col × L*er* reciprocal crosses using *Arabidopsis* lines expressing *PHE1::NTF* and *PHE1::BirA* (lines referred as INT hereafter) [40]. To facilitate the crosses, we used the male sterile mutants *pistillata* (*pi-1*, in L*er* accession) and *dde2* (in Col accession containing INT) as female parents and pollinated them with the INT line (Col accession) and L*er* wild type, respectively. A total of 500mg of siliques for the first replicates and 250mg of siliques for the second and third replicated were collected at 4DAP. Tissue homogenization and nuclei purification were performed from three biological replicates as previously described [40]. Total RNA was extracted from purified nuclei using the mirVana Isolation Kit Protocol (Ambion). mRNA extraction was performed using NEBNext Poly(A) mRNA Magnetic Isolation and the Libraries were prepared with the NEBNext Ultra II RNA Library Prep Kit from Illumina. Samples were sequenced at the National Genomic Infrastructure (NGI) from SciLife Laboratory (Uppsala, Sweden) on an Illumina HiSeq2500 in paired-end 125bp read length. Reads were trimmed and mapped in single-end mode to the Arabidopsis (TAIR10) genome previously masked for *rRNA* genes and for SNP positions between Col and L*er*accessions, using TopHat v2.1. [40] (parameters adjusted as -g 1 -a 10 -i 40 -I 5000 -F 0 -r 130). Gene expression was normalized to reads per kilobase per million mapped reads (RPKM) using GFOLD40 [41]. For discrimination of maternal and paternal transcripts, reads were assigned to the Col or L*er* genomes using single nucleotide polymorphisms (SNPs) between the accessions using SNPsplit [42]. We calculated contamination levels based on the deviation of the read counts from the expected 2:1 maternal/paternal genome ratio in the endosperm [43] (Additional File 1: Table S6). Based on this analysis, the first replicates of both cross directions Col × L*er* and L*er* x Col were not included in the downstream analyses. To increase the statistical power to detect parentally-biased genes, we merged libraries from two replicates of Col × L*er* and three replicates of L*er* × Col for downstream analysis.

### Allele-specific expression analysis

We defined a minimum threshold of 20 informative reads for Col × L*er* (2 replicates) and L*er* × Col (2 replicates) crosses, respectively. Statistical differences between maternal and paternal read counts for each gene were calculated using a Chi-square test, considering genes with a false discovery rate adjusted *P*-value of less than 0.01. Additionally, MEGs required to have at least 85% maternal informative reads in both directions of the reciprocal cross and PEGs to have at least 50% paternal informative reads in both directions of the reciprocal cross, following previously defined conditions [12].

### Generation of reporter constructs and transgenic lines

For the generation of reporter constructs, we used the ClonExpress^®^ MultiS One Step Cloning kit. Promoters (≃ 2kb) and genic sequences of *AT2G33620*, *AT1G43580*, *AT1G47530*, *AT1G64660*, *AT2G30590*, *AT4G15390*, and *AT5G53160* were amplified from WT Col-0 genomic DNA using primers specified in Additional File 1: Table S6 and cloned into vector pB7FWG.0. Constructs were transformed into *Agrobacterium tumefaciens* strain GV3101 and *Arabidopsis* plants were transformed using the floral dip method [44]. Ten transgenic lines per construct were generated and analyzed.

### Imprinting analyses

For reciprocal crosses, designated female partners were emasculated at 1–2 days prior to anthesis. Two days after emasculation pistils were hand-pollinated with respective pollen donors. Seeds were dissected from the siliques and mounted on a microscope slide for imaging and counting at 2 and 4 DAP. For fluorescence analyses, seeds were stained with 0.1 mg/mL propidium iodide (PI) solution in 7% glucose. Seeds of reciprocal crosses of reporter lines were analyzed under confocal microscopy on a Zeiss 800 Inverted Axio Observer with a supersensitive GaASp detector. Images were acquired, analyzed and exported using Zeiss ZEN software. For each reporter construct 50-60 seeds per line were analyzed.

## DECLARATIONS

### Ethics approval and consent to participate

Not applicable

### Consent for publication

Not applicable

### Availability of data and material

The RNA-seq data generated in this study are available through GEO (GSE119915). We furthermore used endosperm RNA expression data from [12], seed coat expression data from [23], parental-specific histone and DNA methylation data from [6], central cell DNA methylation profiles from [3], and DNA methylation data from sperm cell, vegetative cell and endosperm of *fie* and *dme* mutants from [4].

### Authors’ contributions

JMR, VKY and GDTDL executed the experimental procedures. JMR, JSG, GDTDL, VKY and CK analyzed the data. JMR and CK wrote the manuscript. All authors discussed the results and commented on the manuscript.

## Acknowledgements

Sequencing was performed by the SNP&SEQ Technology Platform, Science for Life Laboratory at Uppsala University, a national infrastructure supported by the Swedish Research Council (VRRFI) and the Knut and Alice Wallenberg Foundation.

## Additional files

Additional File 1: **Supplementary Figures**, **Table S3**, **Table S5** and **Table S6**.

Additional File 2: **Table S1.** Scores assigned to genes based on the presence of CHG methylation in the central cell as well as H3K9me2 and H3K27me3 on maternal alleles in the endosperm.

Additional file 3: **Table S2.** Seed coat expression of genes with maximum score (12) based on Additional File 2: Table S1.

Additional file 4: **Table S4.** Parent-of-origin RNAseq dataset of 4DAP INTACT-purified endosperm of Col × L*er* reciprocal crosses.

